# Coastal winds are stronger and impact seabird behaviour and fitness in a warming Arctic

**DOI:** 10.1101/2025.11.19.689241

**Authors:** Tristan J.B. Martin, Jérôme Fort, Andrea S. Grunst, Melissa L. Grunst, Ann M. A. Harding, Françoise Amélineau, Samuel Perret, Fabrice Ardhuin, David Grémillet

## Abstract

The Arctic is warming nearly four times faster than the rest of the planet. This unprecedented warming is profoundly altering environmental conditions. Coastal areas, which are crucial for biodiversity and human activities, are at the forefront of these changes. Among the major environmental changes expected in coastal regions are shifts in wind regimes and an increase in storm frequency. Using an interdisciplinary approach, we demonstrated that (1) wind conditions in Arctic coastal regions during summer are changing, resulting in a rise in strong wind events. (2) These large-scale changes in wind dynamics influence the movements and fitness of seabirds, with broader impacts on panarctic socio-ecosystems. To demonstrate these effects, we used a long-term study of little auks (*Alle alle*), the most abundant Arctic seabird species and an ecological indicator of Arctic coastal ecosystems. We showed that during strong wind events, little auks significantly reduce the visits to the colony. These behavioural changes negatively impact their fitness as inferred from chick growth rates. Overall, strong wind events disrupt the delicate balance between parental investment and self-maintenance in this long-lived species. Over time, such pressures may drive shifts in life-history traits, potentially affecting population dynamics.

## Introduction

The Arctic is warming up to four times faster than the global average (Rantanen et al., 2022), a phenomenon known as Arctic amplification, and mainly driven by changes in albedo, sea ice loss, and oceanic heat transport (Previdi et al., 2021; Serreze & Barry, 2011). As a result, the Arctic undergoes a cascade of abiotic transformations, including increased precipitation, elevated sea surface temperatures, cryosphere declines, intensified hydrological cycles, and coastal erosion (Box et al., 2017; Zang et al., 2023). These changes influence local and global oceanic circulation, triggering feedback loops that affect atmospheric dynamics and enhance extreme weather events (Henderson et al., 2021). Given that approximately 67% of the Arctic region is marine, these climatic shifts have significant implications for aquatic ecosystems: They alter light regimes due to diminishing sea ice, affect ocean stratification, accelerate acidification, increase nutrient fluxes from land to sea, and may disrupt benthic-pelagic coupling (Zhulay et al., 2023; von Appen et al., 2021; Qi et al., 2022). These abiotic transformations are accompanied by profound biological responses. Notably, shifts in environmental conditions impact primary production (Lewis et al., 2020; Grémillet & Descamps, 2023; Clairbaux et al., 2021), with potential cascading effects on Arctic food webs and ecosystem functioning.

Numerous studies have investigated past, ongoing, and projected changes in Arctic systems in response to a variety of climatic and environmental variables (Box et al., 2018; Bintanja and Selten, 2014; Overland et al., 2020; Rantanen et al., 2022; Schuur et al., 2015). Yet, surprisingly, surface wind speed, particularly in coastal areas, has received less attention (Jakobson et al., 2019; Liu et al., 2024; Aperkov et al.,2024). Surface winds play a fundamental role in shaping atmospheric and oceanic dynamics, with far-reaching impacts on ecosystems, human activities, and biophysical processes (Rykaczewski and Checkley, 2008). In the Arctic, coastal wind patterns are of particular importance due to their influence on sea ice circulation, the formation and persistence of polynyas, ocean layer mixing, and the dispersion of nutrients and particles within the water column (Rainville et al., 2011; Jakobson et al., 2019; Du Vivier et al., 2023). These processes are not only critical for maintaining ecological balance, but also have direct implications for Arctic coastal communities and wildlife (Martin et al., 2025). For example, stronger or more variable winds can exacerbate coastal erosion, disrupt traditional fishing and hunting activities, and pose risks to navigation and coastal infrastructures (Ashjian et al., 2010; Barnhart et al., 2014). Similarly, Arctic marine fauna, notably seabirds, rely on predictable wind fields to access feeding grounds and maintain their life cycles (Jakubas et al., 2022). Changes in wind dynamics could disrupt these processes, with cascading effects on food availability and population stability. Wind dynamics are therefore a critical factor influencing seabird ecology (Thorne et al., 2023), particularly for species within the *Alcidae* family (commonly referred to as auks), which represent the most abundant seabird family in the Arctic during summer (Gaston et al., 1998). Auks face unique mechanical constraints because they use their small wings both for flying and diving (Pennycuick, 2008). In auks wings are adapted for flying underwater but, unlike penguins, auks have maintained ability to fly. Consequently, they have some of the highest wing-loading of any bird and flight costs are high (Elliott et al., 2013). Windy conditions can dramatically increase their already high energy expenditure, potentially compromising reproduction and long-term survival (Elliott et al., 2014). Specifically, summer storms can prevent seabirds from flying, and may trigger a “washing-machine effect” (sensu Clairbaux et al. 2021) hindering underwater feeding. The washing-machine effect refers to the disruption of water column stratification, leading to prey dispersion and increased turbidity, which could reduce the foraging efficiency of seabirds (Clairbaux et al. 2021). In addition, strong winds can also degrade the quality of nesting sites. In cliff-nesting auks, for instance, the probability of successful landing decreases as wind speed rises (Shepard et al., 2019), and predation rates by gulls increase under windy conditions (Gilchrist et al., 1998).

Among the auks, the little auk (*Alle alle*) is the smallest species in the Atlantic, and very likely the most abundant Arctic seabird overall, with an estimated population between 40-80 million individuals (Kampp et al., 2000; Isaksen and Gavrilo, 2000; Egevang et al., 2003). Due to its abundance, trophic position, and role in transferring organic matter between marine and terrestrial ecosystems, the little auk is a key ecological engineer and biological indicator in a changing Arctic (Mosbech et al., 2018; González-Bergonzoni et al., 2017). Notably, the species is heavily reliant on zooplankton, whose availability is directly influenced by ocean temperature, sea ice cover and wind conditions (Karnovsky et al. 2010). Little auk fitness therefore reflects the impact of climate change on marine food webs and ecosystem health (Wojczulanis-Jakubas et al., 2022-A). This nutrient transport transforms Arctic barren grounds into vegetated coastal areas, with feedback effects on Arctic climate (González-Bergonzoni et al., 2017).

Recent investigations have demonstrated higher Arctic summer storm frequency in the context of global warming (Zhang et al., 2023) and changes in near surface wind speed at the Arctic scale (Jakobson et al., 2019; Liu et al., 2024). Yet, these analyses did not detail wind conditions within Arctic coastal areas, which are primary seabird foraging grounds in summer. Therefore, our first objective was to analyse panarctic coastal wind between 1979 and 2023, to test the hypothesis that summer wind speeds have increased in recent decades. Second, we used long-term little auk data collected in East Greenland across 2005-2024, to test the hypothesis that summer storms in the Arctic significantly impact seabird foraging performance, with consequences for fitness indicators, notably adult body condition and chick growth rates. We thereby used the notion of fitness in a functional context (Arnold, 1983), with individual proxies of chick growth rate for adult performance and of adult body condition for adult survival propensity (Gebhardt-Henrich and Richner, 1998). More specifically, we predicted negative relationships between wind speed and time spent foraging, number of stormy days and chick growth rates, and number of stormy days and adult body condition at the end of the breeding season that reflect the cumulative impact of strong wind events over the season.

Testing longitudinal trends in coastal summer winds on the Arctic scale is essential for better understanding impacts of climate change on coastal zones, which are home to most Arctic peoples and essential to economically and culturally important wildlife. Also, specifying functional links between wind speed, and the capacity of little auks to acquire food and maintain their fitness, is key to the interpretation and use of little auks as ecological indicators. Overall, understanding the interplay between wind dynamics and seabird fitness is crucial for predicting the resilience of Arctic marine ecosystems to climate change.

## Material & Methods

### Panarctic coastal winds

We analyzed changes in wind speed over Arctic coastal zones within 200 km of the coastline. Wind data were extracted from the European Centre of Medium-range Weather Forecasts (ECMWF) fifth-generation global reanalysis, ERA5 (Hersbach et al., 2020). ERA5 provides hourly weather variables on a global scale, with a spatial resolution of 0.25° × 0.25°. For this study, hourly wind speeds were calculated using the 10m u-component (u₁₀) and v-component of wind (v₁₀), which represent horizontal air speed towards east and north, respectively, at a height of 10 meters above the sea surface. Data for the summer months (June, July, and August) from 1979 to 2023 were used. For each spatial point within coastal areas, the mean and standard deviation (SD) of the hourly wind speed distribution were calculated for each summer. To examine trends in wind mean and SD over time, Sen’s slope was calculated for each Arctic coastal point between 1979 and 2023 (Sen, 1968; Gocic and Trajkovic, 2013). Sen’s slope (*b*) is defined as the median rate of change over time:

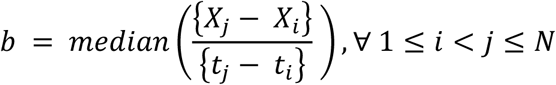

where *X_i_* and *X_j_* are the values of a variable (e.g., mean or SD) at time steps *t_i_* and *t_j_*, respectively, and *N* is the total number of observations in the time series.

Trend significance was evaluated using the Mann–Kendall trend test (Hirsch et al., 1982). Because time series data in the environmental sciences often exhibit temporal autocorrelation, which can affect trend detection, we employed the prewhitening method described by Collaud Coen et al. (2020) to address this issue. Wind trend analyses were performed using Python version 3.12.7 and the *mannkendall* package version 1.1.1 (Coen et al., 2020).

### Seabird data and local weather information

We studied little auks at Ukaleqarteq (Kap Høegh; 70°44’N, 21°35’W) in East Greenland during the breeding season (July–August) from 2005 to 2024. Little auks nest under rocks on steep boulder fields, and raise a single chick per breeding season. After hatching, the chick is brooded by both parents for first few days, then left alone while both parents forage at sea and return several times per day to feed their chick. The chick grows rapidly, reaching around 70% of adult body mass by the time it fledges, which typically occurs 25 days after hatching (Harding et al., 2004). Males and females are similar in size and contribute equally to parental care during linear chick growth (Wojczulanis-Jakubas et al., 2009), so no differentiation between sexes was made.

#### Chick growth

From 2005 to 2007 and 2010 to 2024, chick growth monitoring was conducted annually on 36 chicks on average (Supp. Table 1), from hatching to fledging. One day after hatching and then every two days, measurements to the nearest gram were taken for mass. Chick mass at the end of the breeding season corresponds to the mass of chicks during the mid and late chick-rearing period, i.e., aged 15 days or older (Harding et al., 2004), to integrate wind effects across this crucial part of chick growth. Chick growth rate corresponds to the mass gain divided by the time interval between two measurements and is expressed in g.day^-1^. Chick mass gain is tightly linked to the provisioning capacity of their parents (Hamer et al., 2002).

#### Adult body condition

From 2005 to 2024, adults were caught throughout the breeding season. For each capture, measurements to the nearest gram and millimetre were taken for mass, bill length, right flattened wing length, and right tarsus length. From these data, we calculated the scaled mass index (SMI), based on methods adapted from Peig and Green, (2009) and Grunst et al. (2025), using the formula:

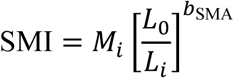

where *L_0_* is the arithmetic mean wing length (a linear size metric), here 120.2mm; Li = length of the linear size metric of individual i; *M_i_* is the measured mass of individual i, and *bSMA* is the coefficient from the regression of natural log (ln) transformed mass against the ln transformed size metric, divided by the correlation coefficient between mass and the size metric (Peig and Green, 2009) here, 2.73. We used flattened wing chord as the linear size metric, as it significantly correlated with body mass (Pearson correlation; r = 0.28, P < 0.05, N = 446). We derived the equation:

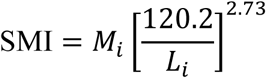

Here, we focused on adult SMI in the mid and late chick-rearing period, when chicks were 15 days or older, on average (Harding et al., 2004).

#### Adult foraging behaviour

At our study site, little auks perform central-place foraging movements between their breeding colony and plankton patches within a range of ca. 100 km from the coast (Amélineau et al. 2016), spending time in-between trips at the colony to feed and broad chicks, and engage in social behaviour with partners and conspecifics (Wojczulanis-Jakubas et al., 2022-B). Between 2017 and 2024, we equipped adult little auks with accelerometers (Axy4, Technosmart; 25 × 10 × 5 mm, 3.0 g including tape for attachment, representing <2% of body mass; see Supp. Table 2 for sample size) to record time activity budgets (TABs) and behavioural traits, such as dive and flight durations. Accelerometers were deployed during the chick-rearing period when chicks were approximately 3–6 days old, based on body size or hatching date. Birds were caught near nests, and accelerometers were attached to ventral feathers using tesa® tape. Handling time was minimized to reduce potential effects on behaviour. Birds were recaptured within 2–11 days.

Acceleration data were recorded in three dimensions at 50 Hz surge (X, back to front), sway (Y, side to side), and heave (Z, dorsal to ventral), while temperature was logged at 0.2 Hz. Data were processed and classified using Igor Pro 9.0 (WaveMetrics, Inc.) with Ethographer 2.05, based on methods adapted from Grunst et al. (2023). Acceleration waves (XW, YW, ZW) were smoothed with a rolling algorithm (1-second sliding window) to extract gravitational acceleration (XW_B_, YW_B_, ZW_B)_. Body pitch was calculated using the formula:

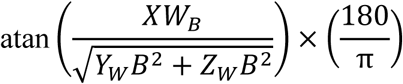

and vectorial dynamic body acceleration (VDBA) was derived as:

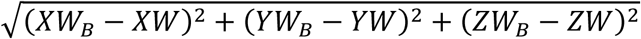

A continuous wavelet transformation was applied to the Z-dimension to estimate dominant wing beat frequency (1 Hz resolution). K-means clustering grouped these signals into clusters for flight (∼10−11 Hz), descending dives (∼3.5−4.5 Hz), resting (∼0 Hz), and intermediate activity (∼0−3.5 Hz), consistent with prior little auk studies (Ste-Marie et al., 2022; Grunst et al., 2023). Behaviours were then classified using clustering outputs, pitch, VDBA, and temperature. Flight was identified by a wing beat frequency of 10−11 Hz. Dives were defined by pitch angles between −10° and −30° with dives ending when pitch rose to >40° before returning to zero. Resting at the colony was identified by the absence of wing beats and an elevated temperature (> 17◦C) following a flight. Resting on water was identified by low wing beat frequency, low temperature (<10◦C), and exclusion from other categories. Through visual inspection, temperature and pitch thresholds were adjusted individually for each bird depending on accelerometer placement. In 2024, thresholds were refined due to minimal differences between air and water temperatures under poor weather conditions. Behavioural classifications were visually validated twice. TABs (flying, resting at the colony, diving, on water) were calculated for the entire accelerometer deployment period. Time spent foraging was calculated as the sum of time spent underwater and in-between dives. As little auks lack circadian rhythms in the Arctic, where there is near 24-hr summer daylight, daily TABs were not analyzed.

### Local weather

For this study, we defined wind speeds ≥ 10 m.s^-1^ as “strong wind”. This threshold corresponds to level 6 on the Beaufort scale, which according to direct observations performed by Inuit hunters, impacts the movements of both birds and coastal human populations (see also: Wang et al., 2024). We used two data sources: from a weather station placed in the colony at 100 meters above sea level on the western slope of the Kap Høegh Peninsula, and from the nearest weather station in Ittoqqortoormiit (40 km away).

At the little auk colony, data were collected at 10 min intervals with a Hobo weather station. Wind speed and direction were measured with a Davis® Wind Speed and Direction Smart Sensor (S-WCF-M003) during the chick-rearing period from 2017 to 2024. We checked for correlations between local weather data collected at our study site and hourly data from the weather station in Ittoqqortoormiit to potentially extend our time series back to 2005 (Cappelen 2019). Unfortunately, the two datasets were poorly correlated. This discrepancy is primarily due to spatial differences causing a time lag of several hours, depending on wind direction. Although the data from the Ittoqqortoormiit weather station do not match the adult foraging behaviour analysis, which requires higher temporal resolution, they still offer a reliable representation of overall weather conditions during the breeding season. In fact, a Spearman correlation test between the number of hours with wind speeds exceeding 10 m.s^-1^ during the chick-rearing period at the fieldwork weather station and the Ittoqqortoormiit weather station yielded a significant result (ρ = 0.94, p < 0.05). Therefore, we used the Ittoqqortoormiit weather data to investigate wind impact on long-term variations in adult body condition and chick mass at the end of the season. We also tested for linkages between data collected at the little auk colony and ERA5 reanalysis data, and encountered no correlation. As a result, for the adult foraging behaviour analysis, we used wind data collected directly at our study site during the last seven years.

### Statistical analyses

Statistical analyses were conducted in R version 4.4.0. To compare the time proportions spent in different behaviours (Resting on water, Dive, Flight, Colony) under two wind conditions (“Low” and “Strong”), we performed non-parametric Wilcoxon rank-sum tests (Using *The stats R package, version 4.4.0*). These tests were performed in accordance with our data structure: individuals observed exclusively under ‘Low’ wind conditions and others observed under both ‘Low’ and ‘Strong’ wind conditions, resulting in unequal sample sizes (n = 34 for Low; n = 13 for Strong).

To examine the relationships between the number of hours of strong wind, adult body condition, and chick weight, we conducted two separate statistical analyses using generalized linear mixed models (GLMMs, using *R package glmmTMB version 1.1.10;* Brooks et al., 2017). These models incorporate random effects to account for variability between years, allowing for a better isolation of wind effects.

To evaluate the impact of wind conditions on chick growth, we tested the influence of the number of hours with strong wind over the two days preceding each chick’s measurement. This two-day interval was chosen in accordance with the two-day sampling rate of the chicks. We used a GLMM (using *R package glmmTMB version 1.1.10;* Brooks et al., 2017) with a Gaussian distribution and an identity link function to investigate factors influencing chick growth rates. We also added chick age as a fixed effect in the model. The model included the year and the sampling day as a random effect. We initially included chick ID as a random effect to account for the dependence between repeated measurements on the same chick and potential parental effects. However, the estimated variance for this effect was close to zero (variance = 9.71×10^-9^ and SD = 9.85×10^-5^), indicating that most of the variation in growth rate was explained by the fixed effects rather than by individual differences between chicks. This suggests that chicks respond homogeneously to environmental conditions. According to AIC values (AIC = 5113.4 and AIC = 5111.4 respectively) and to avoid unnecessary complexity and optimize model fit, we therefore removed this random effect. The model was fit to 1,085 observations, with individuals during 7 breeding seasons. Model significance was determined using z-tests for each fixed effect, with a significance threshold of p < 0.05.

## Results

### Panarctic coastal winds

Panarctic summer near-surface coastal wind speed increased at an average rate of 0.02 m·s⁻¹·yr⁻¹ across 1979-2023, corresponding to an approximate increase of 0.9 m·s⁻¹ (16%) over this period (Figure 1). For mean wind speed, 61.79% of all trends were positive, while 38.21% were negative, regardless of p-value. Among positive trends, 39.30% were statistically significant (p < 0.05), while 26.35% of negative trends were statistically significant (p < 0.05). The standard deviation of summer wind speed also increased at a rate of 0.01 m·s⁻¹·yr⁻¹, corresponding to a ca. 20% increase in wind speed variability over the study period. 67.32% of all trends in wind speed variability were positive, while 32.68% were negative, regardless of statistical significance. Among positive trends, 81.68% were statistically significant (p < 0.05), whereas among negative trends, 59.12% were statistically significant (p < 0.05).

**Figure 1:**
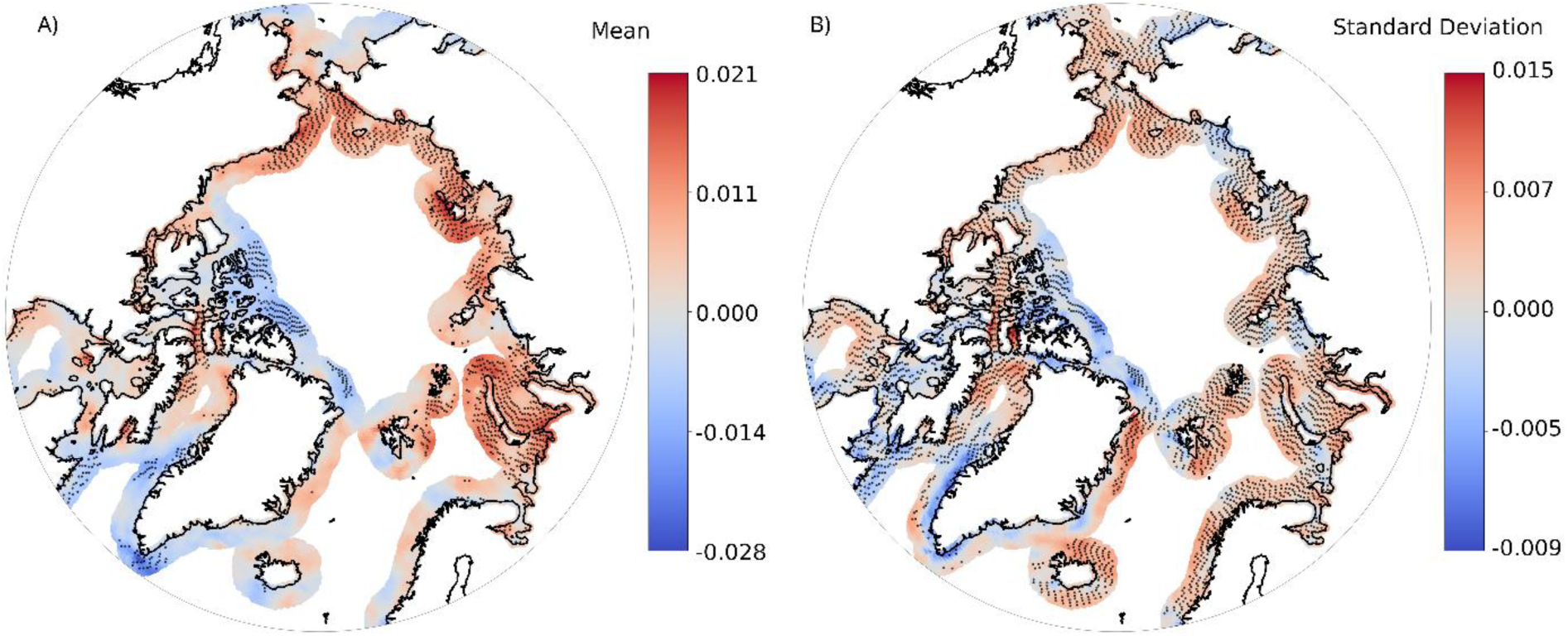
Panarctic summer (June, July, August) near-surface coastal wind speed trends between 1979 and 2023. A) Summer near-surface coastal wind speed mean, and B) summer near-surface coastal wind speed SD. Dots are located where trends are significant.

### Seabird behaviour

Adult little auks exposed to strong winds spent significantly more time foraging (W = 66, p < 0.01), and less time at the colony (W = 417, p < 0.01). Notably, there was a 90% decrease in time spent at the colony during high wind events (Figure 2). Conversely, we found no significant impact of wind conditions on little auk flight time (W = 285, p = 0.132). These findings remained consistent when comparing time budgets within, as well as across individuals exposed to low/high winds.

**Figure 2:**
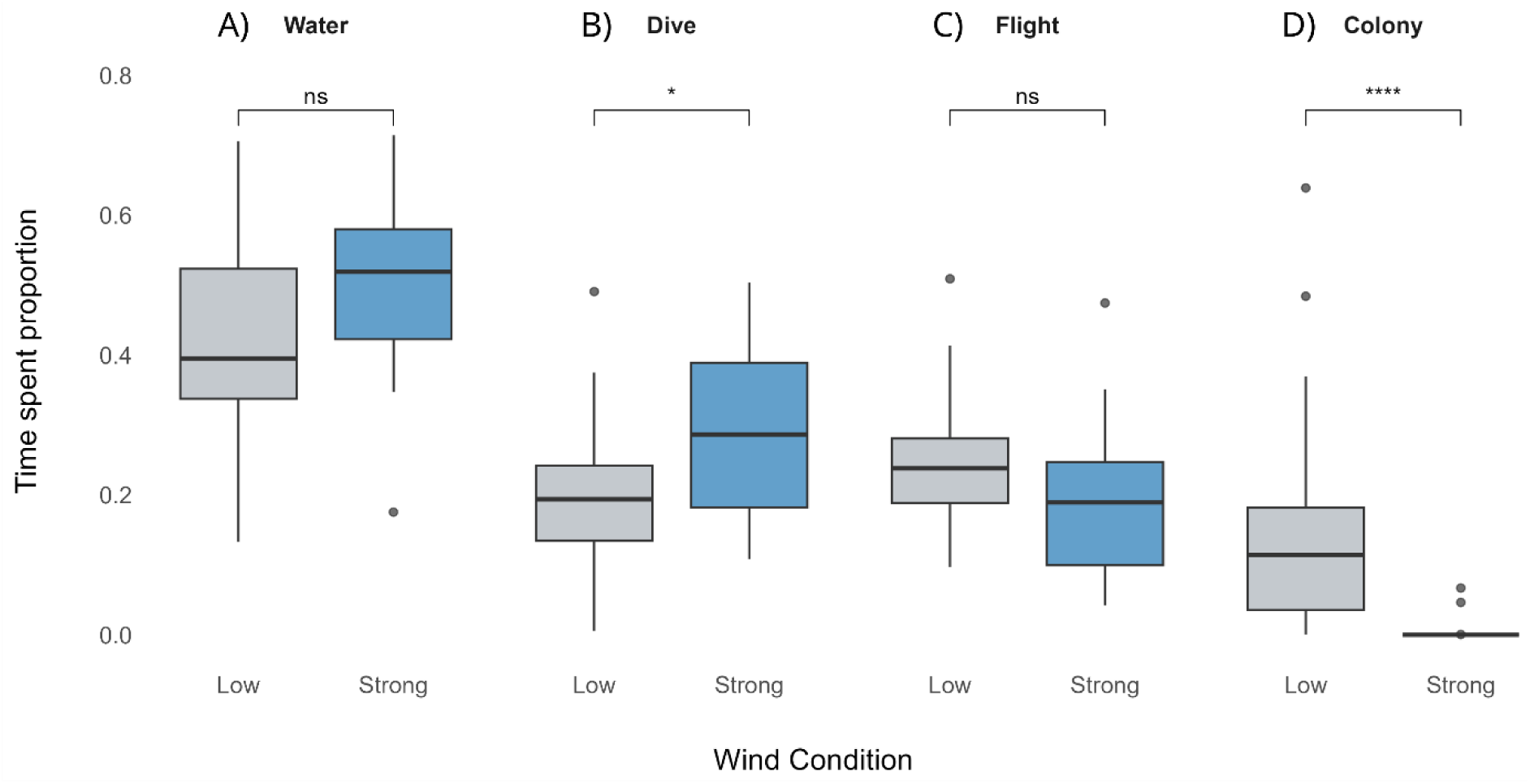
Individual time spent proportion in water, dive, flight and colony behaviour according to wind condition.

### Local weather and seabird fitness

#### a) Adult body condition and chick mass

End-of-season chick weights and adult body conditions were relatively consistent across 2005-2016, but in recent years, there has been greater inter-annual variability. Notably, in 2024, both chick weights (W = 550, p < 0.01) and adult body condition (W = 2272.5, p = 0.046) were abnormally low compare to rest of the time series, a year that also experienced high wind conditions (Figure 3). We further identified a significant relationship (β = −0.36, SE = 0.15, p = 0.015) between the mass of chicks older than 15 days of age and the number of hours of wind during the breeding season. Chick mass was significantly lower as the number of hours of wind during the breeding season increased. However, results were no significant when removing 2024 data (β = −0.15, SE = 0.24, p = 0.53). Also, we did not find any significant relationships between adult body condition at the end of the season and the number of hours of wind during the breeding season (β = −0.11, SE = 0.08, p = 0.15).

**Figure 3:**
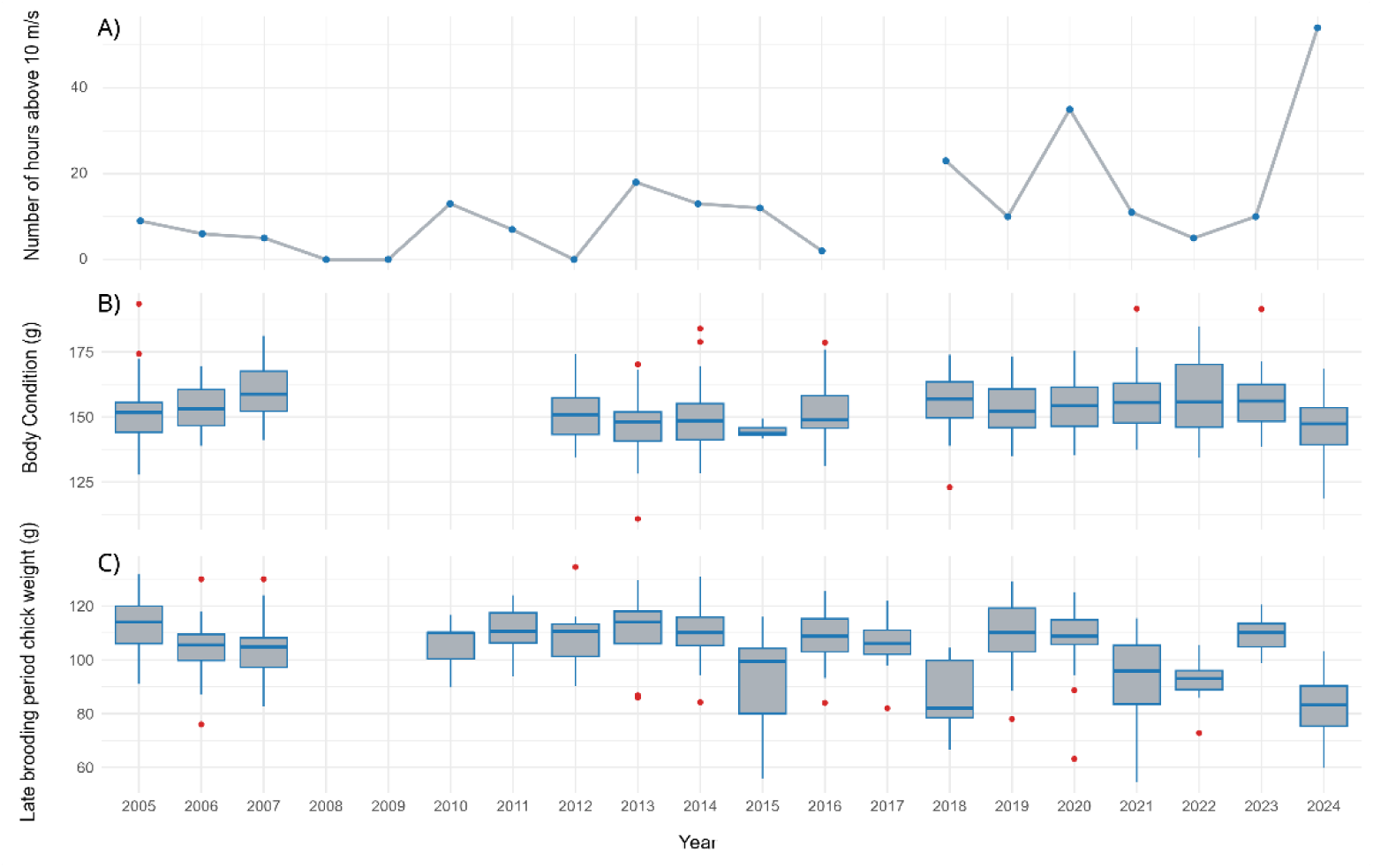
Time series of A-Breeding season number of hours with wind speed ≥ 10 m.s^-1^; B-Mid and late chick-rearing period adult SMI; C-Mid and late chick-rearing period chick weight.

**Figure 4:**
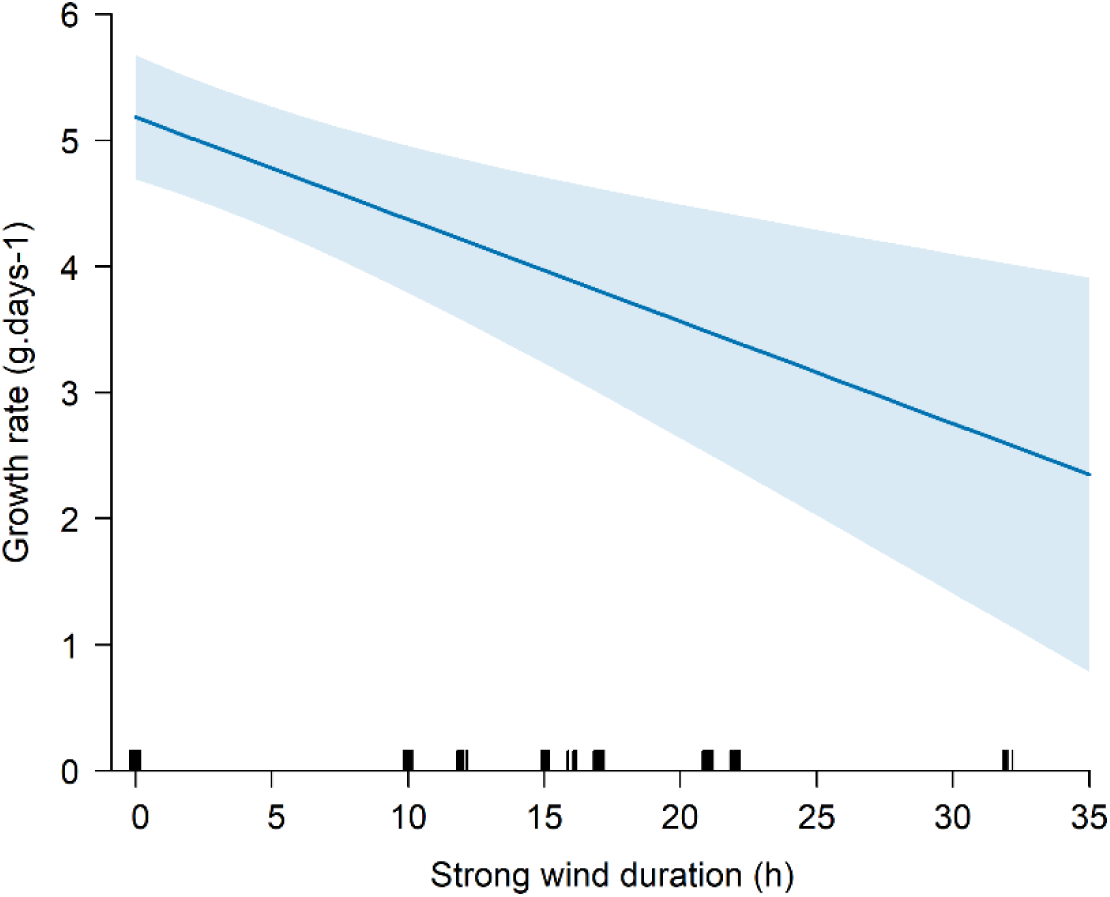
Effect of strong wind duration on chick growth rate and 95% confidence interval, accounting for chick age, and interannual variability.

#### b) Chick growth

The analysis revealed a significant negative relationship between strong wind duration during the two days before measurement and chick growth, with higher strong wind duration associated with lower growth rate (β = −0.081, SE = 0.029, p < 0.01; Figure 3).

## Discussion

Our data clearly demonstrate that climate change over the preceding decades has affected wind conditions during summer in Arctic coastal regions. Since 1979, average summer wind speeds and their variability increased by 16% and 20%, respectively, across most of the Arctic coastline. These changes in windscape at the Arctic scale influence seabird movement, ecology and fitness. Notably, we showed that during strong wind events, little auks remained at sea, with a negative impact on their capacity to provision chicks and fitness, as inferred from chick growth rates and adult body condition toward the end of the linear growth phase.

Our analyses highlight an increase in summer coastal wind speed in the major part of the Arctic in the past 40 years. These results are consistent with other studies that revealed and projected an increase of past and future near surface wind speed at the scale of the Arctic basin (Jakobson et al., 2019; Liu et al., 2024), and reflect broader atmospheric and oceanic changes across the Arctic region (Akperov et al., 2024; DuVivier et al. 2023; Liu et al., 2024; Varvus and Alkama, 2022; Muilwijk et al. 2024). Conversely, strong Arctic coastal winds contrast with the “stilling effect” observed in temperate zones, where wind speeds recently declined due to increased surface roughness and weakened pressure gradients (Vautard et al., 2010). On Arctic scale, wind acceleration is primarily driven by sea ice loss, which reduces surface roughness, enhances turbulent mixing, and facilitates stronger near-surface winds (DuVivier at al. 2023; Liu et al., 2024; Varvus and Alkama, 2022; Muilwijk et al. 2024). Additionally, the diminishing ice cover amplifies ocean-atmosphere temperature contrasts, further intensifying wind speeds (Taylor et al., 2018; Liu et al., 2024; DuVivier at al. 2023). The increase in summer wind variability reflects the rising frequency and persistence of Arctic cyclones, with cyclone tracks invading the Arctic basin in recent decades (DuVivier at al. 2023; Zangh et al., 2023; Semenov et al., 2019). Shifts in atmospheric circulation, including the weakening of the polar vortex and evolving pressure systems, may also play a role in these trends in summer storms and wind conditions. Additionally, recent studies highlight a limitation in the ERA5 reanalysis model, which tends to underestimate wind speeds in coastal regions (Alkhalidi et al., 2025). This bias stems partly from the model’s spatial resolution, which smooths out local variations driven by abrupt topographic gradients and surface roughness transitions at the edges of land and sea ice. In this context, observations indicate a strong correlation between measured and ERA5-predicted wind speeds below 10 m.s^-1^, but a notable underestimation at higher wind speeds, particularly in complex coastal areas with pronounced topographical and thermal contrasts between land and sea (Alkhalidi et al., 2025). Hence, the true extent of ongoing changes in Arctic near surface coastal wind speeds may be underestimated.

High wind events had marked effects on the activity patterns of little auks, with adults significantly reducing the proportion of colony visits. This decrease in colony attendance could be a mechanism designed to avoid elevated predation risk associated with high winds. Indeed, strong winds favor the flight of large bodied predatory birds (Pennycuick, 2008), and the probability of Alcid predation by glaucous gulls (*Larus hyperboreus*) increases during strong winds (Gilchrist and Gaston, 1997). At Brünnich’s guillemot (*Uria lomvia*) colonies, wind provides lift and stability to the glaucous gulls, allowing them to catch guillemots on the nest without landing and confronting their defensive behaviour (Gilchrist et al., 1998). Like guillemots, little auks are prime prey for gulls nesting at the periphery of their colonies. Even though little auks nest under boulders and may hide from gulls, they are, unlike guillemots, too small to fight back, and during strong winds they are particularly vulnerable, both in the air and on the ground.

Low colony attendance during strong winds may also be linked to landing risks. Fast-flying birds’ ability to respond to wind during landing is critical, as errors can lead to injury or even death, likely because wind interferes with the ability to maintain flight control during the final landing phase (Shepard et al., 2019). Finally, little auks may not come back to the colony because of their physical inability to face wind > 17m.s^-1^ (Pennycuick 2008), implying that adults are simply unable to fly and return to land and thus remain at sea during strong wind. This hypothesis is consistent with the fact that little auks spent more time at the water surface during strong winds. They nonetheless maintained similar, although slightly reduced, flight durations, suggesting that they may have been making an effort to return to the colony. This presumption is supported by our direct observations of numerous little auks unsuccessfully battling against strong winds offshore of the colony during storms.

When little auks are stuck at sea during wind gusts, a “washing machine effect”, increases water turbidity, reduces underwater visibility, and scatters prey patches, thereby disrupting little auk ability to gather food (Clairbaux et al., 2021). This is particularly relevant for seabirds such as little auk, which usually forage within the upper 20 meters of the water column during the breeding season. It has also been demonstrated that wind affects the prey distribution of little auks, impacting chick growth rates (Jakubas et al., 2022). In our study, diving behaviour showed a slight increase during periods of strong winds, indicating that the birds persist in their foraging efforts under these conditions. The increase in time spent diving combined with lower chick growth rates supports the idea that strong winds hinder their foraging efficiency. Future studies could integrate GPS and accelerometer data to determine whether little auks remain confined near their colony suggesting constraints linked to predation or landing, or venture further into foraging grounds, thereby supporting the ‘washing machine effect’ hypothesis, wherein strong winds reduce foraging efficiency.

Foragers typically adjust their behaviour to mitigate the effects of environmental conditions on reproduction (Stephens & Krebs 1986). For instance, in Brünnich’s guillemots, wind affects parental behaviour by shifting to alternative food sources on windy days or boosting food delivery rates when conditions improve, but does not influence chick growth rates (Elliot et al., 2014). Yet, when facing unusual stress, long-lived adults are predicted to favor their own survival above that of their offspring (Stearns 2000). This seems to be the case in little auks: previous investigations have demonstrated the species’ capacity to shift diets and increase parental effort when confronted with ocean warming and the disappearance of its preferred foraging habitat, the marginal sea-ice zone (Harding at al., 2009); such major behavioural plasticity allowed birds to maintain both adult body condition and chick growth rates (Grémillet et al. 2012). In great contrast, in our study, we observed a significant decrease in chick growth rates during periods of strong winds, as well as a decrease in chick mass at the end of breeding seasons with numerous and/or long storms. This decline in chick growth is likely to be primarily linked to reduced provisioning by adults during storm events, rather than direct effects of weather conditions on developing nestlings. Indeed, little auk nest below-ground and chicks are usually well-sheltered from the wind. Thus, wind during storm events is unlikely to induce higher thermoregulatory costs.

Adult little auks have displayed admirable resiliency in the face of global change impacts, such as a reduction in the quantity and quality of prey (Wojczulanis-Jakubas et al. 2022). Nevertheless, recent summer storm events threaten to overwhelm the coping mechanisms of this approx. 160g species, which has only a limited capacity to propel itself into the wind. Impacts on fitness through strongly reduced chick growth rate are further underscored by observations performed on 22nd August 2024, one week after little auk peak fledging. prominently affected by storms: Inuit hunters made the unusual encounter of “*thousands of dead little auks*” offshore of our study site in East Greenland. Birds were both adults and fledglings, which tested negative to highly pathogenic avian influenza (Greenlandic Veterinary Authorities, unpubl. data). Their likely cause of death was therefore unusually poor body condition. Such sensitivity to summer storms mirrors the impact of winter cyclones on the survival probabilities of migrating little auks, which may die by the thousand following major winter lows in the North Atlantic (Clairbaux et al. 2021). Our results are also consistent with previous studies on albatrosses which, for instance, show that stronger winds reduce foraging success and increase energetic stress (Darby et al., 2024). Yet, seabird responses vary depending on functional traits, notably wing morphology (Pennycuick 2008; Nourani et al., 2023; Ventura et al., 2024). Strong winds around little auks breeding colonies are also enhanced by a usually steep topography, which favors Venturi effects. Venturi effects correspond to an increase in wind speed that occurs when airflow is funneled through a narrow or constricted space, such as steep topography around breeding colonies. Such habitat is primarily selected by breeders because steep, windswept slopes become snow-free early in the spring and allow little auk access to underground nesting cavities. Yet, such habitat may become a trap as summer wind conditions become increasingly stochastic.

Overall, our study is a major addition to a growing body of research on the impact of extreme climatic events on seabirds. Previous studies have documented consequences of such events, including behavioural responses to storm (Lempidakis et al., 2022), complete breeding failures following snowstorms in Antarctica (Descamps et al. 2015, Descamps et al. 2023), the relocation of nesting sites after intense storms (Bonter et al. 2014), starvation and mass mortalities caused by marine heatwaves (Jones et al. 2018, Piatt et al. 2020, Renner et al., 2024) widespread mortality triggered by cyclones (Lavers et al. 2024), and total reproductive failure due to rainstorms in northern Greenland (Yannick et al. 2014). As seabirds share the coastline with humans, in the Arctic as around the world, wind storms and other extreme climatic events challenge the resilience of seabird communities as well as that of human societies (Martin et al. 2025). It is therefore essential and urgent to understand linkages between climatic conditions and seabird responses, to use the latter as early-warning systems for abrupt transitions into altered ecosystem functioning (Velarde et al., 2019; Solan et al., 2020).

## Acknowledgments

We acknowledge long-term support from the French Polar Institute (IPEV) to the ADACLIM program administered by J. Fort and D. Grémillet. T. Martin is supported by the GDR OMER. We thank Nanu travel for facilitating field seasons, as well as all those involved in data collection. This work also contributes to the SEE-life program funded by CNRS. We are extremely grateful to the many researchers who contributed to the ADACLIM project over two decades. We also thank Greenlandic authorities which gave us permission to perform research, and the people of Ittoqqortoormiit for their support and friendship over the years. Many thanks are due to all colleagues at our respective institutions for their support, in particular to support staff at CNRS as well as at the French Polar Institute IPEV.

## Permits

All field work in East Greenland was conducted in accordance with guidelines for the use of animals (Committee & others, 2024). Experiments were approved by the Danish Polar Center and the Government of Greenland, Ministry of Environment and Nature and Department of Fisheries, Hunting and Agriculture: N° 512–240 (2005), N° 512-258 (2006), N° 07-501 (2007), N° 66.24/23 (2008), N° 66.01.13 (2009 and 2010), N° 2011-047447, N° 2012-065815, N° 2013-083634, N° 2014-098814, N° 2015-115290, N°18789 (2016), N° 2017-13, N° 2018-2257, N° 2019-88, N° 2020-1006, N° 2021-78, N° 2022-196, N° 2023-6417, N° 2024-1943.

## References

Akperov, M., Zhang, W., Koenigk, T., Eliseev, A., Semenov, V. A., & Mokhov, I. I. (2024). Projected changes in near-surface wind speed in the Arctic by a regional climate model. Polar Science, 101162. 10.1016/j.polar.2024.101162

Alkhalidi, M., Al-Dabbous, A., Al-Dabbous, S., & Alzaid, D. (2025). Evaluating the Accuracy of the ERA5 Model in Predicting Wind Speeds Across Coastal and Offshore Regions. Journal of Marine Science and Engineering, 13(1), Article 1. 10.3390/jmse13010149

Amélineau, F., Grémillet, D., Bonnet, D., Bot, T. L., & Fort, J. (2016). Where to Forage in the Absence of Sea Ice? Bathymetry As a Key Factor for an Arctic Seabird. PLOS ONE, 11(7), e0157764. 10.1371/journal.pone.0157764

Arnold, S. J. (1983). Morphology, Performance and Fitness1. American Zoologist, 23(2), 347–361. 10.1093/icb/23.2.347

Ashjian, C. J., Braund, S. R., Campbell, R. G., George, J. C. “Craig,” Kruse, J., Maslowski, W., Moore, S. E., Nicolson, C. R., Okkonen, S. R., Sherr, B. F., Sherr, E. B., & Spitz, Y. H. (2010). Climate Variability, Oceanography, Bowhead Whale Distribution, and Iñupiat Subsistence Whaling near Barrow, Alaska. ARCTIC, 63(2), Article 2. 10.14430/arctic973

Barnhart, K. R., Anderson, R. S., Overeem, I., Wobus, C., Clow, G. D., & Urban, F. E. (2014). Modeling erosion of ice-rich permafrost bluffs along the Alaskan Beaufort Sea coast. Journal of Geophysical Research: Earth Surface, 119(5), 1155–1179. 10.1002/2013JF002845

Bintanja, R., & Selten, F. M. (2014). Future increases in Arctic precipitation linked to local evaporation and sea-ice retreat. Nature, 509(7501), 479–482. 10.1038/nature13259

Bonter, D. N., MacLean, S. A., Shah, S. S., & Moglia, M. C. (2014). Storm-induced shifts in optimal nesting sites: A potential effect of climate change. Journal of Ornithology, 155(3), 631–638. 10.1007/s10336-014-1045-9

Box, J. E., Colgan, W. T., Christensen, T. R., Schmidt, N. M., Lund, M., Parmentier, F.-J. W., Brown, R., Bhatt, U. S., Euskirchen, E. S., Romanovsky, V. E., Walsh, J. E., Overland, J. E., Wang, M., Corell, R. W., Meier, W. N., Wouters, B., Mernild, S., Mård, J., Pawlak, J., & Olsen, M. S. (2019). Key indicators of Arctic climate change: 1971–2017. Environmental Research Letters, 14(4), 045010. 10.1088/1748-9326/aafc1b

Cappelen, J. (n.d.). Denmark – DMI Historical Climate Data Collection 1768-2020.

Clairbaux, M., Mathewson, P., Porter, W., Fort, J., Strøm, H., Moe, B., Fauchald, P., Descamps, S., Helgason, H. H., Bråthen, V. S., Merkel, B., Anker-Nilssen, T., Bringsvor, I. S., Chastel, O., Christensen-Dalsgaard, S., Danielsen, J., Daunt, F., Dehnhard, N., Erikstad, K. E., … Grémillet, D. (2021). North Atlantic winter cyclones starve seabirds. Current Biology, 31(17), 3964–3971.e3. 10.1016/j.cub.2021.06.059

Collaud Coen, M., Andrews, E., Bigi, A., Martucci, G., Romanens, G., Vogt, F. P. A., & Vuilleumier, L. (2020). Effects of the prewhitening method, the time granularity, and the time segmentation on the Mann–Kendall trend detection and the associated Sen’s slope. Atmospheric Measurement Techniques, 13(12), 6945–6964. 10.5194/amt-13-6945-2020

Darby, J., Phillips, R. A., Weimerskirch, H., Wakefield, E. D., Xavier, J. C., Pereira, J. M., & Patrick, S. C. (2024). Strong winds reduce foraging success in albatrosses. Current Biology, 0(0). 10.1016/j.cub.2024.10.018

Descamps, S., Aars, J., Fuglei, E., Kovacs, K. M., Lydersen, C., Pavlova, O., Pedersen, Å. Ø., Ravolainen, V., & Strøm, H. (2017). Climate change impacts on wildlife in a High Arctic archipelago – Svalbard, Norway. Global Change Biology, 23(2), 490–502. 10.1111/gcb.13381

Descamps, S., Hudson, S., Sulich, J., Wakefield, E., Grémillet, D., Carravieri, A., Orskaug, S., & Steen, H. (2023). Extreme snowstorms lead to large-scale seabird breeding failures in Antarctica. Current Biology, 33(5), R176–R177. 10.1016/j.cub.2022.12.055

DuVivier, A. K., Vavrus, S. J., Holland, M. M., Landrum, L., Shields, C. A., & Thaker, R. (2023). Investigating Future Arctic Sea Ice Loss and Near-Surface Wind Speed Changes Related to Surface Roughness Using the Community Earth System Model. Journal of Geophysical Research: Atmospheres, 128(20), e2023JD038824. 10.1029/2023JD038824

Egevang, C., Boertmann, D., Mosbech, A., & Tamstorf, M. P. (2003). Estimating colony area and population size of little auks Alle alle at Northumberland Island using aerial images. Polar Biology, 26(1), 8–13. 10.1007/s00300-002-0448-x

Elliott, K. H., Chivers, L. S., Bessey, L., Gaston, A. J., Hatch, S. A., Kato, A., Osborne, O., Ropert-Coudert, Y., Speakman, J. R., & Hare, J. F. (2014). Windscapes shape seabird instantaneous energy costs but adult behavior buffers impact on offspring. Movement Ecology, 2(1), 17. 10.1186/s40462-014-0017-2

Gaston, A. J., Jones, I. L., & Lewington, I. (1998). The auks: Alcidae. Bird Families of the World. https://www.waterbouwkundiglaboratorium.be/nl/catalogus

Gilchrist, H. G., & Gaston, A. J. (1997). Effects of murre nest site characteristics and wind conditions on predation by glaucous gulls. Canadian Journal of Zoology, 75(4), 518–524. 10.1139/z97-064

Gilchrist, H. G., Gaston, A. J., & Smith, J. N. M. (1998). Wind and Prey Nest Sites as Foraging Constraints on an Avian Predator, the Glaucous Gull. Ecology, 79(7), 2403–2414. 10.1890/0012-9658(1998)079[2403:WAPNSA]2.0.CO;2

Gebhardt-Henrich, S., & Richner, H. (1998). Causes of growth variation and its consequences for fitness. Oxford Ornithology Series, 8, 324–339.

Gocic, M., & Trajkovic, S. (2013). Analysis of changes in meteorological variables using Mann-Kendall and Sen’s slope estimator statistical tests in Serbia. Global and Planetary Change, 100, 172–182. 10.1016/j.gloplacha.2012.10.014

González-Bergonzoni, I., Johansen, K. L., Mosbech, A., Landkildehus, F., Jeppesen, E., & Davidson, T. A. (2017). Small birds, big effects: The little auk (Alle alle) transforms high Arctic ecosystems. Proceedings of the Royal Society B: Biological Sciences, 284(1849), 20162572. 10.1098/rspb.2016.2572

Grémillet, D., & Descamps, S. (2023). Ecological impacts of climate change on Arctic marine megafauna. Trends in Ecology & Evolution, 38(8), 773–783. 10.1016/j.tree.2023.04.002

Grémillet, D., Welcker, J., Karnovsky, N. J., Walkusz, W., Hall, M. E., Fort, J., Brown, Z. W., Speakman, J. R., & Harding, A. M. A. (2012). Little auks buffer the impact of current Arctic climate change. Marine Ecology Progress Series, 454, 197–206. 10.3354/meps09590

Grunst, A. S., Grunst, M. L., Grémillet, D., Kato, A., Bustamante, P., Albert, C., Brisson-Curadeau, É., Clairbaux, M., Cruz-Flores, M., Gentès, S., Perret, S., Ste-Marie, E., Wojczulanis-Jakubas, K., & Fort, J. (2023). Mercury Contamination Challenges the Behavioral Response of a Keystone Species to Arctic Climate Change. Environmental Science & Technology, 57(5), 2054–2063. 10.1021/acs.est.2c08893

Grunst, M. L., Grunst, A., Grémillet, D., Amiguet, M., Charrier, J., Cruz-Flores, M., Lacoue-Labarthe, T., & Fort, J. (2025). Arctic Seabirds Exposed to Acute Stress Display State- and Environment-Dependent Patterns of Surface Temperature Change, Independent of Mercury Contamination (SSRN Scholarly Paper 5122660). Social Science Research Network. 10.2139/ssrn.5122660

Hamer, K. C., Schreiber, E. A., & Burger, J. (2002). Breeding biology, life histories, and life history-environment interactions in seabirds. Biology of marine birds, 217–261.

Harding, A. M. A., Kitaysky, A. S., Hall, M. E., Welcker, J., Karnovsky, N. J., Talbot, S. L., Hamer, K. C., & Grémillet, D. (2009). Flexibility in the parental effort of an Arctic-breeding seabird. Functional Ecology, 23(2), 348–358. 10.1111/j.1365-2435.2008.01488.x

Harding, A. M. A., Van Pelt, T. I., Lifjeld, J. T., & Mehlum, F. (2004). Sex differences in Little Auk Alle alle parental care: Transition from biparental to paternal-only care. Ibis, 146(4), 642–651. 10.1111/j.1474-919X.2004.00297.x

Henderson, G. R., Barrett, B. S., Wachowicz, L. J., Mattingly, K. S., Preece, J. R., & Mote, T. L. (2021). Local and Remote Atmospheric Circulation Drivers of Arctic Change: A Review. Frontiers in Earth Science, 9. 10.3389/feart.2021.709896

Hersbach, H., Bell, B., Berrisford, P., Hirahara, S., Horányi, A., Muñoz-Sabater, J., Nicolas, J., Peubey, C., Radu, R., Schepers, D., Simmons, A., Soci, C., Abdalla, S., Abellan, X., Balsamo, G., Bechtold, P., Biavati, G., Bidlot, J., Bonavita, M., … Thépaut, J.-N. (2020). The ERA5 global reanalysis. Quarterly Journal of the Royal Meteorological Society, 146(730), 1999–2049. 10.1002/qj.3803

Hirsch, R. M., Slack, J. R., & Smith, R. A. (1982). Techniques of trend analysis for monthly water quality data. Water Resources Research, 18(1), 107–121. 10.1029/WR018i001p00107

Isaksen, K. & Gavrilo, M. (2000) Little auk Alle alle. In The status of marine birds breeding in the Barents Sea region (ed. T. Anker-Nilssen, V. Bakken, H. Strøm, A. N. Golovkin, V. V. Bianki and I. P. Tatarinkova), pp 131-136. Rapport No 113. Oslo: Norsk Polarinstitutt.

Jakobson, L., Vihma, T., & Jakobson, E. (2019). Relationships between Sea Ice Concentration and Wind Speed over the Arctic Ocean during 1979–2015. 10.1175/JCLI-D-19-0271.1

Jakubas, D., Wojczulanis-Jakubas, K., Szeligowska, M., Darecki, M., Boehnke, R., Balazy, K., Trudnowska, E., Kidawa, D., Grissot, A., Descamps, S., & Błachowiak-Samołyk, K. (2022). Gone with the wind – Wind speed affects prey accessibility for a High Arctic zooplanktivorous seabird, the little auk Alle alle. Science of The Total Environment, 852, 158533. 10.1016/j.scitotenv.2022.158533

Jones, T., Parrish, J. K., Peterson, W. T., Bjorkstedt, E. P., Bond, N. A., Ballance, L. T., Bowes, V., Hipfner, J. M., Burgess, H. K., Dolliver, J. E., Lindquist, K., Lindsey, J., Nevins, H. M., Robertson, R. R., Roletto, J., Wilson, L., Joyce, T., & Harvey, J. (2018). Massive Mortality of a Planktivorous Seabird in Response to a Marine Heatwave. Geophysical Research Letters, 45(7), 3193–3202. 10.1002/2017GL076164

Kampp, K., Falk, K., & Egevang Pedersen, C. (2000). Breeding density and population of little auks (Alle alle) in a Northwest Greenland colony. Polar Biology, 23(8), 517–521. 10.1007/s003000000115

Karnovsky, N., Harding, A., Walkusz, W., Kwaśniewski, S., Goszczko, I., Jr, J. W., Routti, H., Bailey, A., McFadden, L., Brown, Z., Beaugrand, G., & Grémillet, D. (2010). Foraging distributions of little auks Alle alle across the Greenland Sea: Implications of present and future Arctic climate change. Marine Ecology Progress Series, 415, 283–293. 10.3354/meps08749

Lavers, J. L., Mead, T. M., Fidler, A. L., & Bond, A. L. (2024). Cyclone Ilsa in April 2023 led to significant seabird mortality on Bedout Island. Communications Earth & Environment, 5(1), 1–9. 10.1038/s43247-024-01342-6

Lempidakis, E., Shepard, E. L. C., Ross, A. N., Matsumoto, S., Koyama, S., Takeuchi, I., & Yoda, K. (2022). Pelagic seabirds reduce risk by flying into the eye of the storm. Proceedings of the National Academy of Sciences, 119(41), e2212925119. 10.1073/pnas.2212925119

Lewis, K. M., van Dijken, G. L., & Arrigo, K. R. (2020). Changes in phytoplankton concentration now drive increased Arctic Ocean primary production. Science, 369(6500), 198–202. 10.1126/science.aay8380

Liu, W., Yang, S., Chen, D., Zha, J., Zhang, G., Zhang, Z., Zhang, T., Xu, L., Hu, X., & Deng, K. (2024). Rapid Acceleration of Arctic Near-Surface Wind Speed in a Warming Climate. Geophysical Research Letters, 51(8), e2024GL109385. 10.1029/2024GL109385

Martin, T., Henri, D. A., Martinez, L.-M., Chandelier, M., Ardhuin, F., & Grémillet, D. (2025). A common future for coastal peoples and seabirds facing extreme climatic events. ICES Journal of Marine Science, 82(1), fsae198. 10.1093/icesjms/fsae198

Mateos, M., & Arroyo, G. M. (2011). Ocean surface winds drive local-scale movements within long-distance migrations of seabirds. Marine Biology, 158(2), 329–339. 10.1007/s00227-010-1561-y

Mosbech, A., Johansen, K. L., Davidson, T. A., Appelt, M., Grønnow, B., Cuyler, C., Lyngs, P., & Flora, J. (2018). On the crucial importance of a small bird: The ecosystem services of the little auk (Alle alle) population in Northwest Greenland in a long-term perspective. Ambio, 47(2), 226–243. 10.1007/s13280-018-1035-x

Muilwijk, M., Hattermann, T., Martin, T., & Granskog, M. A. (2024). Future sea ice weakening amplifies wind-driven trends in surface stress and Arctic Ocean spin-up. Nature Communications, 15(1), 6889. 10.1038/s41467-024-50874-0

Nourani, E., Safi, K., Grissac, S. de, Anderson, D. J., Cole, N. C., Fell, A., Grémillet, D., Lempidakis, E., Lerma, M., McKee, J. L., Pichegru, L., Provost, P., Rattenborg, N. C., Ryan, P. G., Santos, C. D., Schoombie, S., Tatayah, V., Weimerskirch, H., Wikelski, M., & Shepard, E. L. C. (2023). Seabird morphology determines operational wind speeds, tolerable maxima, and responses to extremes. Current Biology, 33(6), 1179–1184.e3. 10.1016/j.cub.2023.01.068

Overland, J., Dunlea, E., Box, J. E., Corell, R., Forsius, M., Kattsov, V., Olsen, M. S., Pawlak, J., Reiersen, L.-O., & Wang, M. (2019). The urgency of Arctic change. Polar Science, 21, 6–13. 10.1016/j.polar.2018.11.008

Peig, J., & Green, A. J. (2009). New perspectives for estimating body condition from mass/length data: The scaled mass index as an alternative method. Oikos, 118(12), 1883–1891. 10.1111/j.1600-0706.2009.17643.x

Pennycuick, C. J. (2008). Modelling the Flying Bird. Elsevier.

Piatt, J. F., Parrish, J. K., Renner, H. M., Schoen, S. K., Jones, T. T., Arimitsu, M. L., Kuletz, K. J., Bodenstein, B., García-Reyes, M., Duerr, R. S., Corcoran, R. M., Kaler, R. S. A., McChesney, G. J., Golightly, R. T., Coletti, H. A., Suryan, R. M., Burgess, H. K., Lindsey, J., Lindquist, K., … Sydeman, W. J. (2020). Extreme mortality and reproductive failure of common murres resulting from the northeast Pacific marine heatwave of 2014-2016. PLOS ONE, 15(1), e0226087. 10.1371/journal.pone.0226087

Previdi, M., Smith, K. L., & Polvani, L. M. (2021). Arctic amplification of climate change: A review of underlying mechanisms. Environmental Research Letters, 16(9), 093003. 10.1088/1748-9326/ac1c29

Qi, D., Ouyang, Z., Chen, L., Wu, Y., Lei, R., Chen, B., Feely, R. A., Anderson, L. G., Zhong, W., Lin, H., Polukhin, A., Zhang, Y., Zhang, Y., Bi, H., Lin, X., Luo, Y., Zhuang, Y., He, J., Chen, J., & Cai, W.-J. (2022). Climate change drives rapid decadal acidification in the Arctic Ocean from 1994 to 2020. Science, 377(6614), 1544–1550. 10.1126/science.abo0383

Rainville, L., Lee, C. M., & Woodgate, R. A. (2011). Impact of Wind-Driven Mixing in the Arctic Ocean. Oceanography, 24(3), 136–145.

Rantanen, M., Karpechko, A. Yu., Lipponen, A., Nordling, K., Hyvärinen, O., Ruosteenoja, K., Vihma, T., & Laaksonen, A. (2022). The Arctic has warmed nearly four times faster than the globe since 1979. Communications Earth & Environment, 3(1), 168. 10.1038/s43247-022-00498-3

Renner, H. M., Piatt, J. F., Renner, M., Drummond, B. A., Laufenberg, J. S., & Parrish, J. K. (2024). Catastrophic and persistent loss of common murres after a marine heatwave. Science, 386(6727), 1272–1276. 10.1126/science.adq4330

Rykaczewski, R. R., & Checkley, D. M. (2008). Influence of ocean winds on the pelagic ecosystem in upwelling regions. Proceedings of the National Academy of Sciences, 105(6), 1965–1970. 10.1073/pnas.0711777105

Ste-Marie, E., Grémillet, D., Fort, J., Patterson, A., Brisson-Curadeau, É., Clairbaux, M., Perret, S., Speakman, J. R., & Elliott, K. H. (2022). Accelerating animal energetics: High dive costs in a small seabird disrupt the dynamic body acceleration–energy expenditure relationship. Journal of Experimental Biology, 225(12), jeb243252. 10.1242/jeb.243252

Schuur, E. a. G., McGuire, A. D., Schädel, C., Grosse, G., Harden, J. W., Hayes, D. J., Hugelius, G., Koven, C. D., Kuhry, P., Lawrence, D. M., Natali, S. M., Olefeldt, D., Romanovsky, V. E., Schaefer, K., Turetsky, M. R., Treat, C. C., & Vonk, J. E. (2015). Climate change and the permafrost carbon feedback. Nature, 520(7546), 171–179. 10.1038/nature14338

Semenov, A., Zhang, X., Rinke, A., Dorn, W., & Dethloff, K. (2019). Arctic Intense Summer Storms and Their Impacts on Sea Ice—A Regional Climate Modeling Study. Atmosphere, 10(4), Article 4. 10.3390/atmos10040218

Sen, P. K. (1968). Estimates of the Regression Coefficient Based on Kendall’s Tau. Journal of the American Statistical Association, 63(324), 1379–1389. 10.2307/2285891

Serreze, M. C., & Barry, R. G. (2011). Processes and impacts of Arctic amplification: A research synthesis. Global and Planetary Change, 77(1), 85–96. 10.1016/j.gloplacha.2011.03.004

Shepard, E., Cole, E.-L., Neate, A., Lempidakis, E., & Ross, A. (2019). Wind prevents cliff-breeding birds from accessing nests through loss of flight control. eLife, 8, e43842. 10.7554/eLife.43842

Solan, M., Archambault, P., Renaud, P. E., & März, C. (2020). The changing Arctic Ocean: Consequences for biological communities, biogeochemical processes and ecosystem functioning. Philosophical Transactions of the Royal Society A: Mathematical, Physical and Engineering Sciences, 378(2181), 20200266. 10.1098/rsta.2020.0266

Stearns, S. C. (2000). Life history evolution: Successes, limitations, and prospects. Naturwissenschaften, 87(11), 476–486. 10.1007/s001140050763

Stephens, D. W., & Krebs, J. R. (1986). Foraging Theory. Princeton University Press.

Taylor, P. C., Hegyi, B. M., Boeke, R. C., & Boisvert, L. N. (2018). On the Increasing Importance of Air-Sea Exchanges in a Thawing Arctic: A Review. Atmosphere, 9(2), Article 2. 10.3390/atmos9020041

Thorne, L., Clay, T., Phillips, R., Silvers, L., & Wakefield, E. (2023). Effects of wind on the movement, behavior, energetics, and life history of seabirds. Marine Ecology Progress Series, 723, 73–117. 10.3354/meps14417

Vautard, R., Cattiaux, J., Yiou, P., Thépaut, J.-N., & Ciais, P. (2010). Northern Hemisphere atmospheric stilling partly attributed to an increase in surface roughness. Nature Geoscience, 3(11), 756–761. 10.1038/ngeo979

Vavrus, S. J., & Alkama, R. (2022). Future trends of arctic surface wind speeds and their relationship with sea ice in CMIP5 climate model simulations. Climate Dynamics, 59(5), 1833–1848. 10.1007/s00382-021-06071-6

Velarde, E., Anderson, D. W., & Ezcurra, E. (2019). Seabird clues to ecosystem health. Science, 365(6449), 116–117. 10.1126/science.aaw9999

Ventura, F., Sander, N., Catry, P., Wakefield, E., Pascalis, F. D., Richardson, P. L., Granadeiro, J. P., Silva, M. C., & Ummenhofer, C. C. (2024). Oceanic seabirds chase tropical cyclones. Current Biology, 34(14), 3279–3285.e3. 10.1016/j.cub.2024.06.022

von Appen, W.-J., Waite, A. M., Bergmann, M., Bienhold, C., Boebel, O., Bracher, A., Cisewski, B., Hagemann, J., Hoppema, M., Iversen, M. H., Konrad, C., Krumpen, T., Lochthofen, N., Metfies, K., Niehoff, B., Nöthig, E.-M., Purser, A., Salter, I., Schaber, M., … Boetius, A. (2021). Sea-ice derived meltwater stratification slows the biological carbon pump: Results from continuous observations. Nature Communications, 12, 7309. 10.1038/s41467-021-26943-z

Wang, K., Guo, Y., Wu, D., Zheng, C., & Wu, K. (2024). Arctic Wind, Sea Ice, and the Corresponding Characteristic Relationship. Journal of Marine Science and Engineering, 12(9), Article 9. 10.3390/jmse12091511

Wojczulanis-Jakubas, K., Jakubas, D., Kidawa, D., & Kośmicka, A. (2012). Is the transition from biparental to male-only care in a monogamous seabird related to changes in body mass and stress level? Journal of Ornithology, 153(3), 793–800. 10.1007/s10336-011-0796-9

Wojczulanis-Jakubas, K., Jakubas, D., & Stempniewicz, L. (2022-A). The Little Auk Alle alle: An ecological indicator of a changing Arctic and a model organism. Polar Biology, 45(2), 163–176. 10.1007/s00300-021-02981-7

Wojczulanis-Jakubas, K., Grissot, A., Devogel, M., Altmeyer, L., Fujisaki, T., Jakubas, D., Kidawa, D., & Karnovsky, N. (2022-B). Post-foraging in-colony behaviour of a central-place foraging seabird. Scientific Reports, 12(1), 12981. 10.1038/s41598-022-17307-8

Yannic, G., Aebischer, A., Sabard, B., & Gilg, O. (2014). Complete breeding failures in ivory gull following unusual rainy storms in North Greenland. Polar Research, 33(1), 22749. 10.3402/polar.v33.22749

Zhang, X., Tang, H., Zhang, J., Walsh, J. E., Roesler, E. L., Hillman, B., Ballinger, T. J., & Weijer, W. (2023). Arctic cyclones have become more intense and longer-lived over the past seven decades. Communications Earth & Environment, 4(1), 348. 10.1038/s43247-023-01003-0

Zhulay, I., Iken, K., Renaud, P. E., Kosobokova, K., & Bluhm, B. A. (2023). Reduced efficiency of pelagic–benthic coupling in the Arctic deep sea during lower ice cover. Scientific Reports, 13(1), 6739. 10.1038/s41598-023-33854-0

